## Supplementary data for "Integrative modelling of the full-length human dehydrodolichyl diphosphate synthase using a hybrid computational and experimental approach"

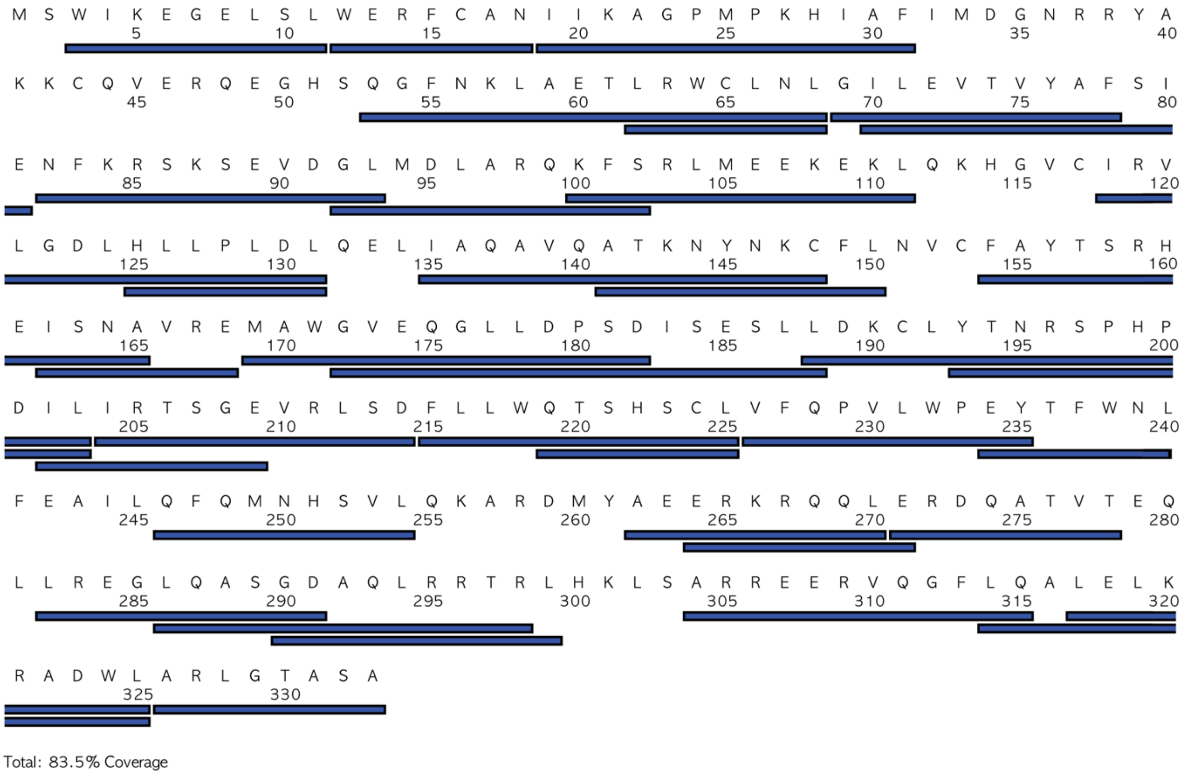


**Figure S1. HDX-MS analysis of human DHDDS.** Sequence coverage of human DHDDS.

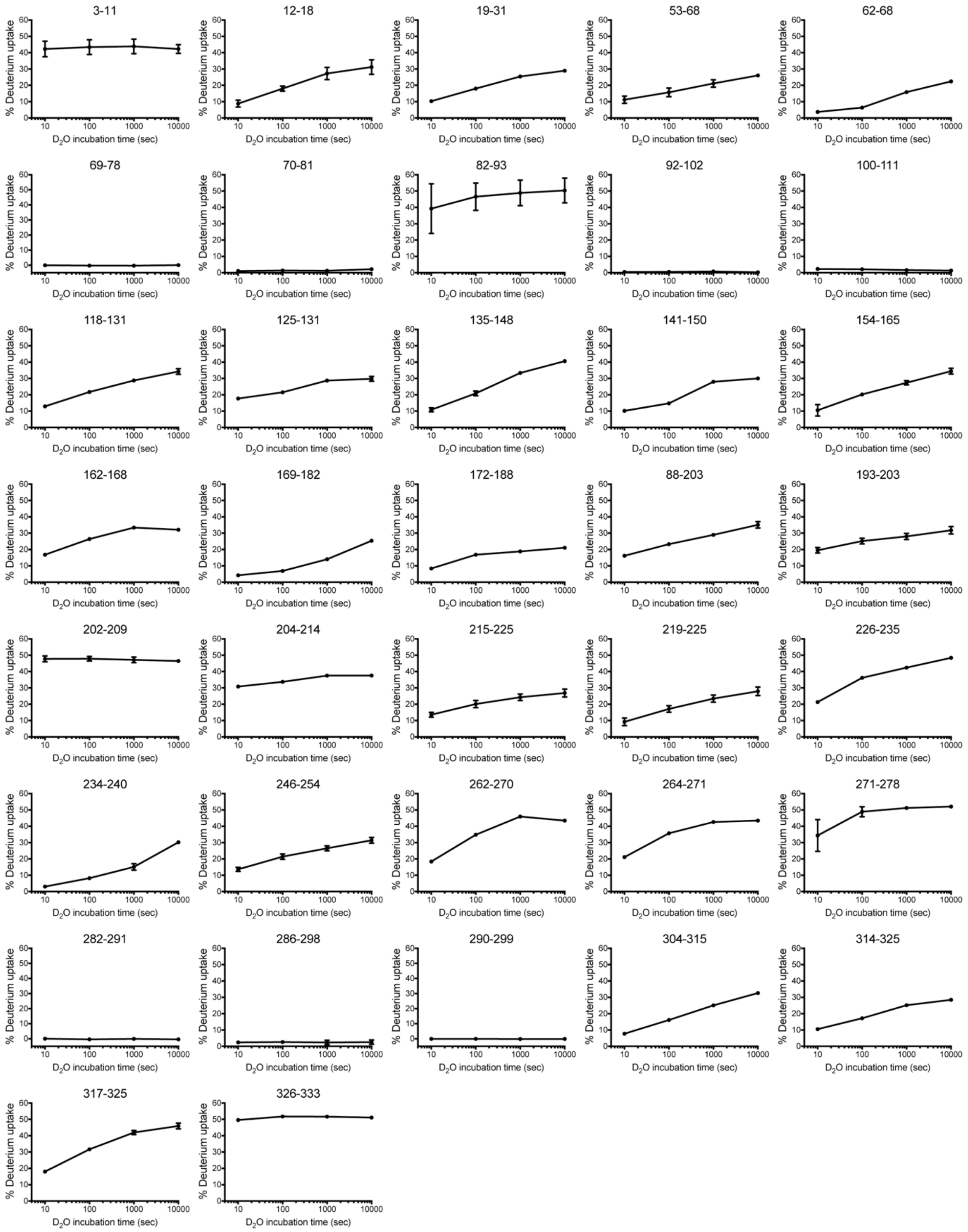


**Figure S2. HDX-MS analysis of human DHDDS.** Deuterium uptake plots.
